# ROS-mediated TNFR Wengen activation in response to apoptosis

**DOI:** 10.1101/2023.11.13.566843

**Authors:** José Esteban-Collado, Mar Fernàndez-Mañas, Manuel Fernández-Moreno, Ignacio Maeso, Montserrat Corominas, Florenci Serras

## Abstract

The activation of tumor necrosis factor receptors (TNFR) controls pleiotropic pro-inflammatory functions ranging from apoptosis to survival. The ability to trigger a particular function will depend on the upstream activation, association with regulatory complexes and downstream pathways. In *Drosophila,* two TNFRs have been identified, Wengen (Wgn) and Grindelwald (Grnd). Although several reports associate these receptors with JNK-dependent apoptosis, it has recently been found that Wgn activates a variety of functions. We demonstrate that Wgn is required for survival by protecting cells from apoptosis. This is mediated by the signaling molecule dTRAF1 and results in the activation of the p38 MAP kinase signaling pathway. Remarkably, Wgn is required for apoptosis-induced regeneration and is activated by the reactive oxygen species (ROS) produced following apoptosis. This ROS activation is exclusive for Wgn, but not for Grnd, and occurs in the absence of the ligand Eiger/TNFα. Furthermore, based on protein sequence conservation, the extracellular Cys-rich domain of Grnd is much more divergent and phylogenetically restricted than that of Wgn, which is more similar to TNFR families from other animals, including those of human TNFRs. Taken together, our results show a novel function for a TNFR that responds to cellular damage by ensuring the cell survival required for the response to damage.

## Introduction

The cytokine Tumor Necrosis Factor α (TNFα) is rapidly released after trauma, injury or infection and acts as a central mediator in the inflammatory response. It signals through the TNF receptors (TNFRs) and its function is currently understood to be pleiotropic, playing a role in apoptosis, cell survival and cell proliferation, the outcome of which will be determined by different TNFRs and complex signaling networks (Gough and Myles, 2020).

Oxidative stress, generated by the production of reactive oxygen species (ROS), is recognized as an underlying cause of a variety of inflammatory diseases (Conner & Grisham, 1996; Droge, 2002; Finkel, 2011). ROS produce deleterious effects on cells because of their ability to oxidize cell structures, such as DNA (Van Houten et al., 2018), membrane lipids and proteins (Colquhoun, 2010; Moriarty-Craige & Jones, 2004). However, it is increasingly well-accepted that ROS-induced post-translational modifications of proteins may also be of physiological relevance in cell signaling. Among these is the oxidation of cysteine residues in receptor proteins, which results in their activation (Lipton et al., 2002; Truong & Carroll, 2012). Likewise, oxidative stress can modify the reduced thiol groups of the TNFR1 extracellular cysteine-rich domain (CRD), which is the hallmark domain shared by the TNFR superfamily (Dominici et al., 2004). This raises the intriguing possibility that oxidative stress might modulate TNFR signaling. Indeed, oxidative stress promotes self-association of the subunits of mammalian TNFR1 and 2, which results in ligand-independent signaling as well as enhanced ligand-dependent TNF signaling (Ozsoy et al., 2008).

Tolerable levels of ROS can propagate as paracrine signals and modulate the intracellular machinery that will reconstruct the damaged tissues (Serras, 2022). In the presence of ROS, thioredoxin dissociates from the MAP3 kinase Apoptosis signal-regulating kinase-1 (Ask1), following which Ask1 oligomerizes, autophosphorylates, recruits its partners and forms an active kinase complex (Bunkoczi et al., 2007; Matsuzawa, 2016; Obsil & Obsilova, 2017; Takeda et al., 2008). An active Ask1 catalytic domain triggers the phosphorylation of the MAP2 kinases that in turn phosphorylate JNK and p38 to induce apoptosis (Bunkoczi et al., 2007; Ichijo et al., 1997; Tobiume et al., 2001). But Ask1 has other functions beyond apoptosis. In *Drosophila*, Ask1 in the gut and imaginal epithelia operates as a survival signal in tissue regeneration by triggering a non-apoptotic function of p38 (Patel et al., 2019; Santabárbara-Ruiz et al., 2019). This is achieved by a ROS-dependent, Akt-induced phosphorylation of Ser83 in Ask1 (Ser174 in humans) that is essential to redirect the Ask1 signaling pathway towards p38 and thus drive regeneration (Esteban-Collado et al., 2021).

The kinase activity of Ask1 is also stimulated by TNFα via members of the TNF Receptor-Associated Factor (TRAF) family of adaptor proteins (Ichijo et al., 1997; Nishitoh et al., 1998). TNFα can transduce the TRAF-mediated signal from TNFR to Ask1 to modulate its activity (Hoeflich et al., 1999; Nishitoh et al., 1998, p. 0; Obsil & Obsilova, 2017; Shiizaki et al., 2013). How this TNF/TNFR/TRAF axis drives apoptosis or survival is poorly understood. In contrast with the large families of both TNF and TNFRs in mammals, *Drosophila* has only one TNFα ortholog, *eiger* (*egr*), and two TNFRs, *wengen* (*wgn*) and *grindelwald* (*grnd*). Grnd mediates the pro-apoptotic function of Egr/TNFα, and its overexpression activates JNK-dependent apoptosis (Andersen et al., 2015; Palmerini et al., 2021). Wengen was first discovered as a TNFR that is able to bind to TRAF, trigger JNK-dependent apoptosis, and transduce signals from Egr/TNFα (Geuking et al., 2005; Kanda et al., 2002; Kauppila et al., 2003). However, it has been shown that while the knockdown of *grnd* blocks apoptosis, that of *wgn* does not (Andersen et al., 2015), suggesting that *wgn* and *grnd* are not redundant. Moreover, both TNFRs are transmembrane proteins that form hexamers for ligand-joining, but Wgn binds to Egr/TNFα at an affinity that is three orders of magnitude lower than Grnd (Palmerini et al., 2021). Recent work has demonstrated that in the gut, Wgn suppresses dTRAF3-mediated lipolysis independently of its ligand and maintains tissue homeostasis, while Wgn-dTRAF2-mediated suppression of immunity occurs in an Egr-dependent manner (Loudhaief et al., 2023). In addition, the ligand-independent function of Wgn has been recently demonstrated to associate with unrelated factors such as FGFR to regulate vesicle trafficking during tracheal development (Letizia et al., 2023). Wgn is also expressed in photoreceptor progenitors and binds to moesin for axonal pathfinding in a ligand-independent manner (Ruan et al., 2013). Moreover, *Drosophila* TNFRs show different subcellular localization. Grnd is mainly found on the apical side of epithelial cells and becomes internalized upon binding to Egr, whereas Wgn is mainly found in intracellular vesicles (Andersen et al., 2015; Letizia et al., 2023; Loudhaief et al., 2023). So far, the evolutionary origin of the two *Drosophila* TNFRs and their relationship to TNFRs from other species have not been investigated in detail, limiting our understanding of how the contrasting functions and molecular behavior of these two receptors may have originated.

Therefore, we used the wing imaginal disc epithelium to explore whether these *Drosophila* TNFRs are involved in survival or apoptosis, whether they respond to oxidative stress and, ultimately, whether they are required for apoptosis-induced regeneration. Here we show that Wgn/dTRAF1 is required for cell survival, in contrast to the apoptotic role of Grnd/dTRAF2. Evolutionary analyses of TNFRs showed an independent and ancient origin of *grnd* and *wgn*, reinforcing the idea of a subfunctionalization of these genes and a higher degree of similarity between the CRDs of Wgn and the TNFRs of humans and other deuterostomes. Indeed, Wgn, but not Grnd, is required for regeneration and for the activation of p38 in the damage response. Interestingly, the activation of Wgn is Egr/TNFα-independent and is sensitive to the ROS produced by damaged cells.

## Results

### Wgn is required for survival by protecting cells from apoptosis

We first studied the involvement of the TNFRs Grnd and Wgn in apoptosis and survival. The two *Drosophila* TNFRs are normally expressed in the wing imaginal discs (Palmerini et al., 2021), whereas Eiger/TNFα (*egr*) is not. We ectopically expressed *egr* in the wing disc, using the *Gal4/UAS/Gal80^TS^*transactivation system, and simultaneously knocked down the TNFRs using appropriate RNAi strains. We used the *hedgehog-Gal4* strain (hereafter *hh>*) to activate the transcription of *egr* in the posterior compartment using the *UAS-egr^weak^* transgene (hereafter *egr^weak^*), a strain that produces a reduced Egr/TNFα activity (Moreno et al., 2002).

The expression of *egr^weak^* resulted in low levels of apoptotic cells in the posterior compartment of the disc (Fig. 1A). However, *egr^weak^* expression resulted in a strong abolition of apoptosis when *grnd* was downregulated (Fig. 1B), which coincides with previous findings that Grnd promotes JNK-dependent apoptosis (Andersen et al., 2015; Palmerini et al., 2021). In contrast, *egr^weak^* expression resulted in a strong enhancement of apoptosis when *wgn* was downregulated (Fig. 1E). This observation was corroborated with two independent *RNAi-wgn* strains (Fig. S1A and S1B).

**Figure 1.**
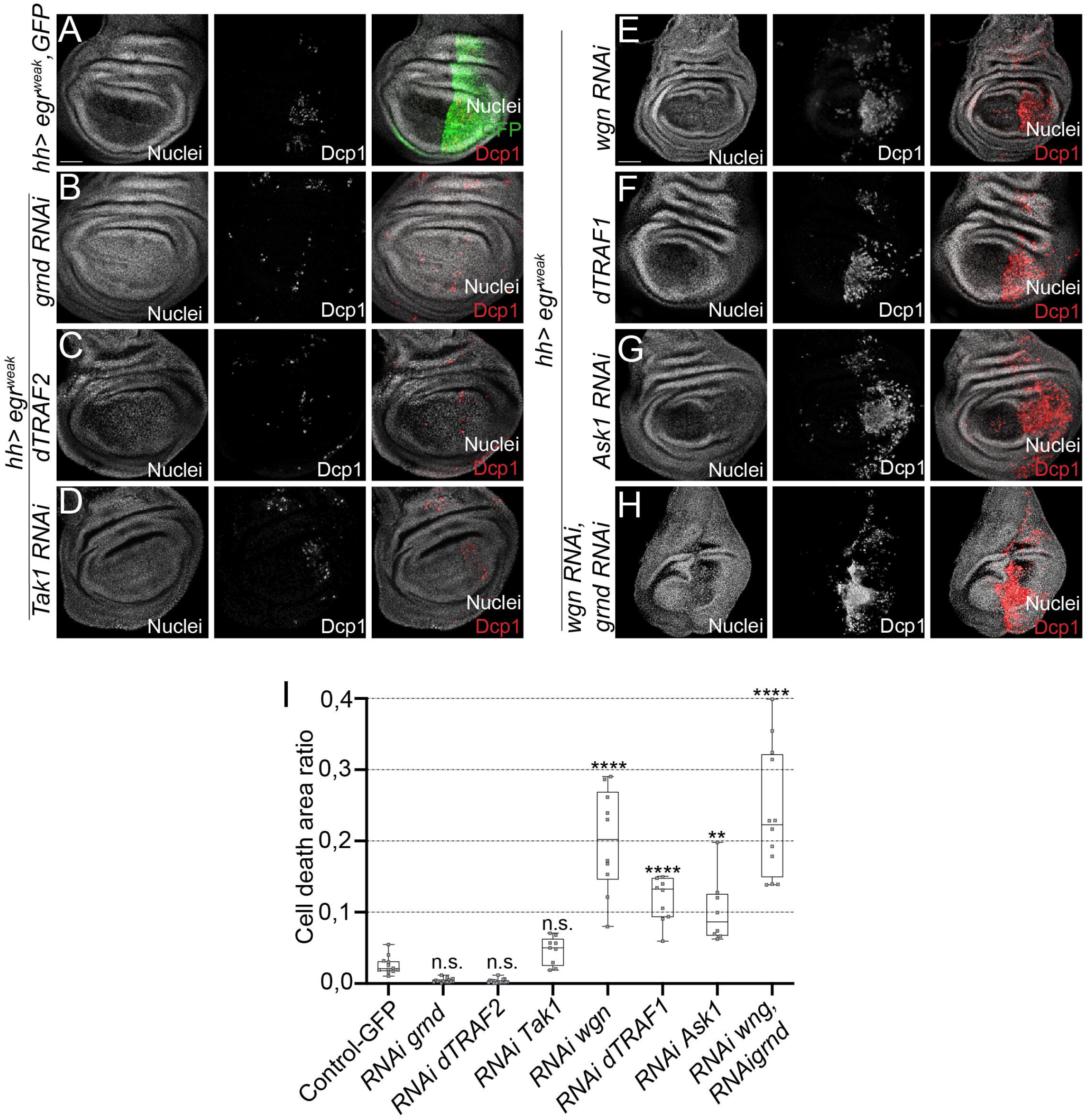
Induction of apoptosis in Egr/TNFα cells after downregulation of different TNFR pathway genes. (A-H) Dcp1-positive cells (red) after ectopic *egr^weak^*expression in (A) GFP-expressing control (n=12), (B) *RNAi grnd* (n=15), (C) *RNAi dTRAF2* (n=18), (D) *RNAi Tak1* (n=9), (E) *RNAi wgn* (n=18), (F) *RNAi dTRAF1*(n=10), (G) *RNAi Ask1* (n=8) and (H) double RNAi of *wgn* and *grnd* (n=12). (I) Cell death area ratio calculated in base 1 for the genotypes indicated. Box plots show maximum-minimum range (whiskers), upper and lower quartiles (open rectangles), and median value (horizontal black line), each dot representing the cell death area ratio from a different imaginal disc. n.s.=not significant, \*\**p*=0.0002, \*\*\*\**p*<0.0001. TP3 used to stain nuclei. Scale bar: 50µm.

Next, we knocked down the adaptor protein *dTRAF2* and the MAP3 kinase *Tak1* and found reduced or similar levels of apoptosis than the *egr^weak^* alone, respectively (Fig. 1C, 1D, 1I) confirming the role of those genes in mediating the pro-apoptotic function of TNF signaling. In contrast, knocking down *dTRAF1* and *Ask1* (Fig. 1F-G) resulted in increased apoptosis in comparison to *egr^weak^* alone (Fig. 1F, 1G, 1I).

To check for epistasis between *grnd* and *wgn*, we activated *hh> egr^weak^* and knocked down both TNFRs. We found high levels of cell death compared to *wgn RNAi* alone (Fig. 1H and 1I), which demonstrates that *wgn* down-regulation is dominant over *grnd*.

These results show that *wgn* is required in cell survival, likely through the Wgn/DTRAF1/Ask1 signaling cassette. In addition, as *grnd* has been demonstrated to be involved in JNK-dependent apoptosis (Andersen et al., 2015), our observations also indicate that the functions of *wgn* and *grnd* are essentially different.

### Wgn and Grnd are phylogenetically divergent

To track the differences observed between *grnd* and *wgn*, we decided to investigate the evolutionary origin of these two *Drosophila* genes. This involved searching for homologs in other animal lineages and performing maximum likelihood (ML) phylogenetic analyses using the amino acid sequences of the CRDs, the only conserved protein domain shared by all TNFRs (Fig. 2A and S2A). We observed that *Drosophila* Grnd and Wgn CRD sequences were in separate branches of the tree, in two different, highly supported monophyletic groups that together included most of the TNFRs genes we had identified in arthropods (Supplementary Table 1). We also assigned the few remaining arthropod TNFRs to either Grnd or Wgn groups through an additional phylogenetic analysis using the full-length protein sequences (see Fig. S2B and Methods). The Wgn branch included genes from multiple arthropod groups, ranging from dipterans such as *Drosophila*, to chelicerates, indicating that Wgn has a very ancient origin that predates the diversification of crown arthropods before the Cambrian, ∼546 million years ago (mya) (Lozano-Fernandez et al., 2020) (Fig. 2B). Grnd had a more restricted taxonomic distribution and was only present in pancrustacean species (insects plus crustaceans), suggesting that this gene family had a slightly younger origin that was nevertheless extremely ancient (emerging at least 514 mya) (Wolfe et al., 2016) (Fig. 2B). Thus, these results showed that Grnd and Wgn are very different from an evolutionary perspective, suggesting they originated through an ancient duplication event of an ancestral TNFR in pancrustaceans followed by a highly asymmetric evolution (Holland et al., 2017) or even that there is an independent origin of the *grnd* and *wgn* CRDs (i.e. through independent recruitment/exon shuffling with cysteine-rich regions from other gene families). Consistent with this, Grnd and Wgn clades were more closely related to some of the TNFR gene families from deuterostome species than they were to each other (Fig. 2C and Fig. S2C). However, in this case the bootstrap supports were too low to reliably assign orthology relationships, probably because CRD sequences are too divergent and short (less than 60 aa) to robustly establish evolutionary relationships between TNFR genes from distant animal phyla (i.e. human and insects), as has been previously suggested (Huang et al., 2008). Therefore, we applied an alternative approach to assess similarities between the deuterostome TNFRs, Grnd, and Wgn and performed a PCA-based alignment (Fig. 2C) and a pairwise BLAST comparison (Fig. S2C). In both analyses, Grnd CRD sequences clustered separate from all other TNFRs, while Wgn family members exhibited a higher degree of similarity with deuterostome TNFRs, clustering together with some of the CRD sequences from hemichordates (the acorn worm, *Saccoglossus kowalevskii*), non-vertebrate chordates (the European amphioxus, *Branchiostoma lanceolatum*) and vertebrates (*Homo sapiens*). Thus, Grnd would constitute a more divergent member of the TNFR superfamily than Wgn, a notion also supported by the unique and derived pattern of cysteine residues that characterize Grnd CRDs, which contain 8 conserved cysteines instead of the 6 cysteines typically found in most TNFR families (Fig. S2D) (Palmerini et al., 2021). These results reinforce the hypothesis of the repurposing of *Drosophila grnd* and *wgn* for different biological functions. The expansion and subfunctionalization of TNFR for apoptosis, survival, and the stress response has been suggested for other metazoan clades (Quistad & Traylor-Knowles, 2016).

**Figure 2.**
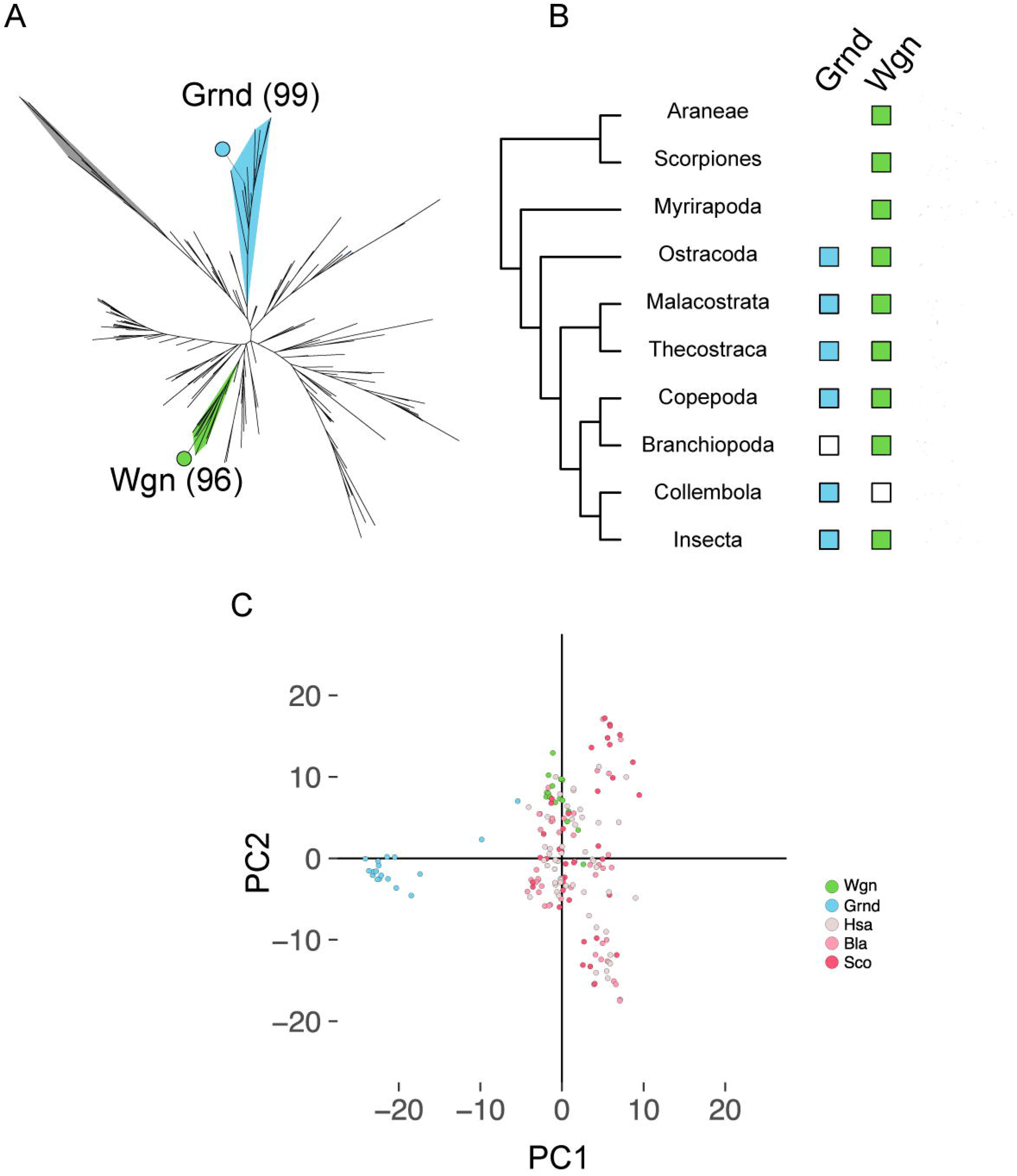
Evolution of Wgn and Grnd. (A) ML phylogenetic tree of the CRDs of Wgn, Grnd, and deuterostome TNFRs. *Drosophila melanogaster* Wgn and Grnd are indicated by solid green and light blue circles, and their corresponding monophyletic clades are colored accordingly. The outgroup branch, containing the cysteine-rich sequences of LIM proteins, is indicated in gray. Wgn and Grnd presence/absence in different arthropod lineages. The presence of the two gene families is indicated by solid squares with the same colors as in (A); white squares indicate putative gene losses. Divergence times of crown arthropods and crown pancrustaceans are indicated at the corresponding nodes of the arthropod tree. (C) PCA-based alignment of CRDs of Wgn, Grnd, and Deuterostome TNFRs.

### The protective role of Wgn is mediated by the activation of the p38 MAP kinase

Next, we analyzed the function of Wgn in cell survival. Although Ask1 can activate JNK and p38 (Tobiume et al., 2001), stimulation of p38 and inhibition of the JNK-proapoptotic function is necessary for a regenerative response to damage (Patel et al., 2019; Santabárbara-Ruiz et al., 2019). Therefore, we hypothesized that the survival role driven by Wgn is likely accomplished by p38 rather than JNK.

To explore this hypothesis, we tested if the apoptosis induced by knocking down *wgn* can be abolished by ectopic stimulation of JNK or p38. We used *hh>egr^weak^, RNAi wgn,* which exacerbate apoptosis, and expressed the MAP2 kinases (*UAS-hep^wt^* or *UAS-lic^wt^*) upstream of either JNK or p38. An ectopic activation of the JNK pathway by *UAS-hep^wt^* resulted in increased levels of apoptosis in comparison to the control (*RNAi wgn* alone) (Fig. 3A, 3C, 3D). In contrast, an ectopic activation of p38 by *lic^wt^* resulted in a strong decrease in apoptosis (Fig. 3A, 3B, 3D), suggesting that activated p38 functions downstream of Wgn for cell survival.

**Figure 3.**
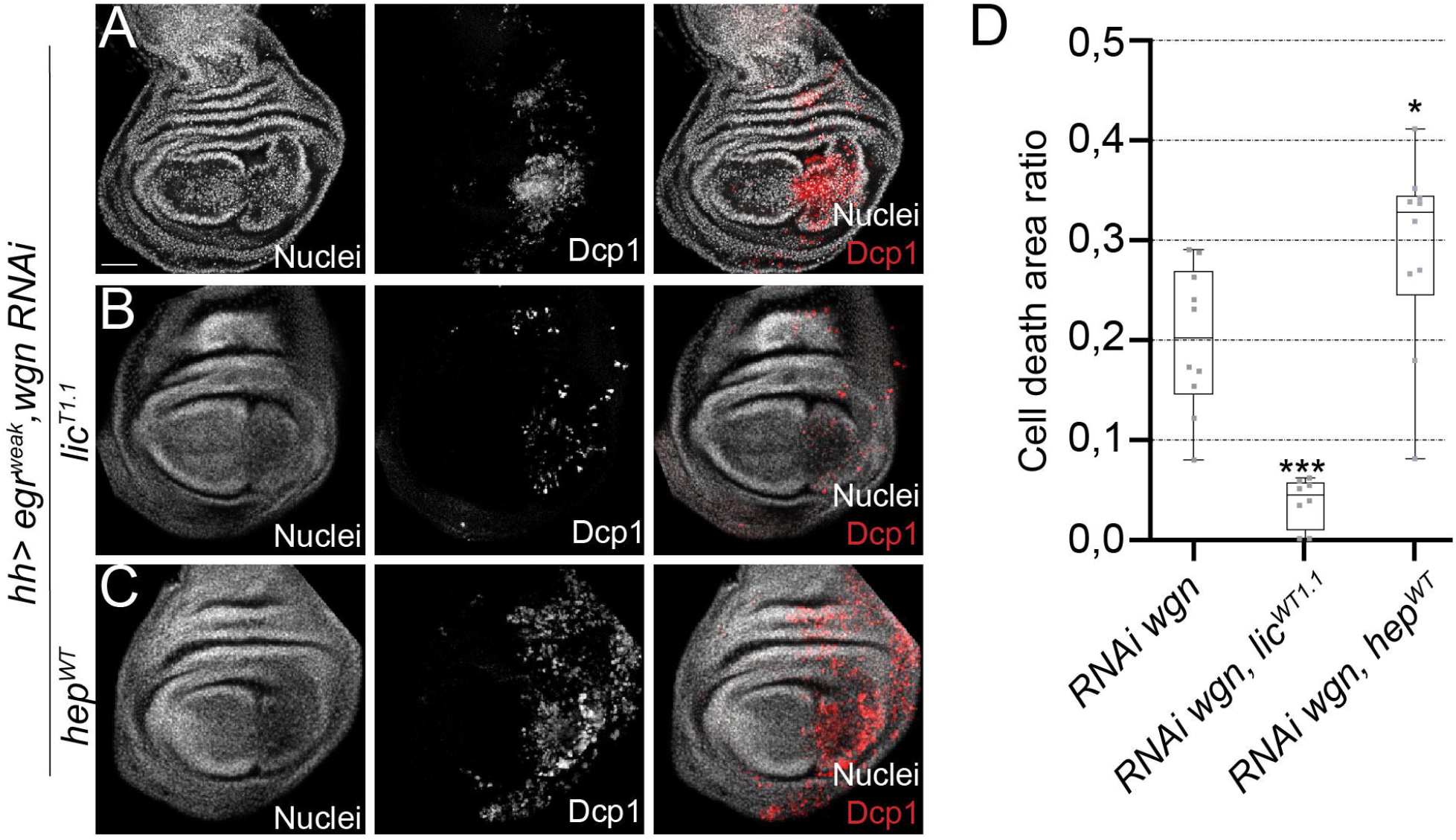
The p38 MAP2 kinase *lic* rescues the *wgn* mutant phenotype. (A-C) Dcp1-positive cells after *egr^weak^* overexpression and *RNAi wgn* in (A) *RNAi wgn* (n=10), (B) *RNAi wgn* and *lic^wt^* (n=8), (C) *RNAi wgn* and *hep^wt^* (n=10). (E) Cell death area ratio calculated in base 1, for the genotypes indicated. Box plots show maximum-minimum range (whiskers), upper and lower quartiles (open rectangles), and median value (horizontal black line). \**p*=0.0314, \*\*\**p*=0.0002. TP3 used to stain nuclei. Scale bar: 50µm.

### Wgn is activated by ROS

Next, we investigated whether Wgn responds to the stress generated by the expression of *egr*. To this end, we first examined the localization of Wgn and Grnd after *egr^weak^* expression. It has been reported that the majority of Wgn is localized in the cytoplasm in many organs, likely in intracellular vesicles rather than at the plasma membrane (Letizia et al., 2023; Loudhaief et al., 2023; Palmerini et al., 2021). By contrast, Grnd is localized in the plasma membrane, making it more accessible to Egr/TNFα (Palmerini et al., 2021). We confirmed these localization patterns for Wgn and Grnd in control imaginal discs (Fig. 4A and 4B).

**Figure 4.**
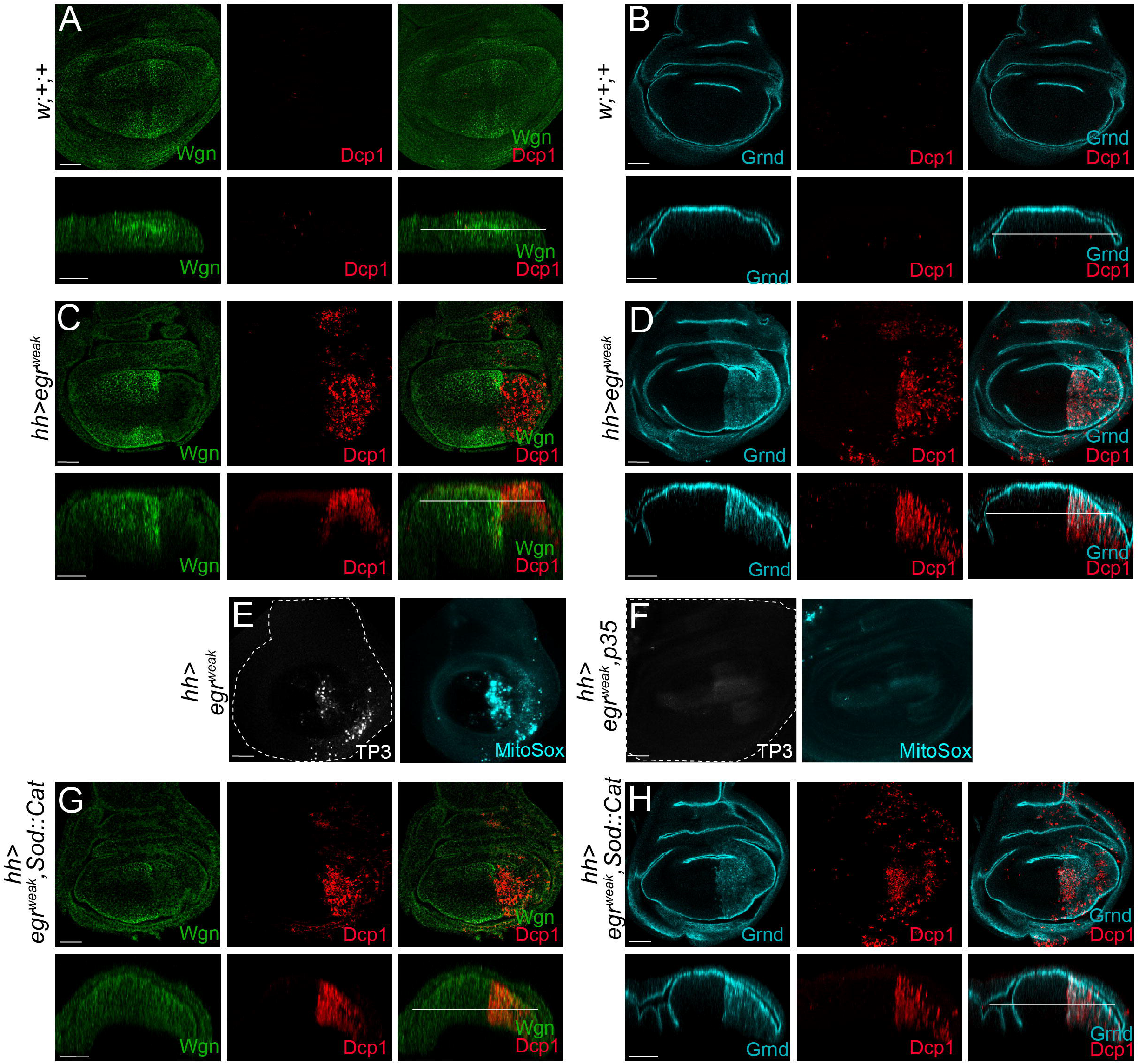
Wgn accumulation after ectopic *egr/TNF*α expression is abrogated after enzymatic depletion of ROS. All transgenes were ectopically expressed in the posterior compartment (*hh>*). The wing pouch of each genotype (above) is complemented with an orthogonal apico-basal section (below); white lines indicate the level of the z-axis of the above images. (A) Anti-Wgn and Dcp1 staining of control wing discs (n=11). (B) Anti-Grnd and Dcp1 staining of control wing discs (n=4). (C) Anti-Wgn and Dcp1 staining of *egr^weak^* (n=10). (D) Anti-Grnd and Dcp1 staining of *egr^weak^*(n=7). (E-F) Discs stained ex-vivo after *egr^weak^* activation. Left: cell death (TP3, white). Right: ROS of mitochondrial origin (MitoSOX, cyan) in (E) *RNAi wgn* and (F) *RNAi wgn* and ectopic expression of p35; dotted lines indicate the contour of the imaginal discs. (G) Anti-Wgn and Dcp1 staining of *egr^weak^* and *Sod1*:*Cat* (n=8). (H) Anti-Grnd and Dcp1 staining of *egr^weak^* and *Sod1*:*Cat* (n=8). Scale bars: 50µm.

After ectopic *egr^weak^* expression in the posterior compartment, Wgn particles were found accumulated in the cytoplasm of the anterior compartment, particularly in anterior cells close to the border with the posterior compartment, and they were absent in the *egr^weak^* posterior compartment (Fig. 4C). By contrast, Grnd was maintained on the apical membrane of the anterior compartment and was found internalized in the *egr^weak^* posterior compartment (Fig. 4D), which agrees with previous observations that Grnd is translocated from the membrane after binding to Egr/TNFα (Andersen et al., 2015).

It is known that apoptosis generates oxidative stress due to the production of ROS of mitochondrial origin that can propagate to recruit neighboring cells for damage repair (Serras, 2022). Furthermore, the oxidation of the TNFR CRD by ROS is a physiological mechanism able to transduce the signal (Ozsoy et al., 2008). Thus, to investigate the molecular mechanism underlying the accumulation of Wgn in neighboring cells after *egr* expression, we first checked if ROS were produced after *hh>egr^weak^*. We used in vivo imaging with the cell-permeant fluorogenic probe MitoSOX, which is non-fluorescent in the reduced state and exhibits bright fluorescence upon oxidation. MitoSOX-positive cells were found in the *egr^weak^* compartment and they co-localized with cells positive for TO-PRO-3, a stain that is very sensitive to dead and dying cells (Fig. 4E). To discern whether ROS production responds to *egr^weak^* expression or to apoptosis, we expressed the baculovirus protein p35, which blocks the effector caspases (Hay & de Belleroche, 1994). We found that neither MitoSOX nor TO-PRO-3 were detected when apoptosis was blocked in *egr^weak^*-expressing cells (Fig. 4F). This suggests that apoptosis causes the accumulation of ROS rather than *egr* expression.

Next, we enzymatically decreased ROS production using ectopic expression of the ROS scavengers *Superoxide dismutase 1* and *Catalase* (*UAS-Sod1:UAS-Cat*). This resulted in a reduction of the accumulation of Wgn particles near the ablated area compared to *egr^weak^* alone (Fig. 4G). In contrast, Grnd localization was not altered after ROS depletion (Fig. 4H).

As the expression of the *egr^weak^* results in apoptosis, we decided to induce apoptosis in an Egr-independent manner to monitor the localization of both TNFRs. With this aim, we expressed the pro-apoptotic gene *reaper (*rpr) using the *sal^E/Pv^-Gal4* driver, whose expression is restricted to the central part of the wing imaginal disc (henceforth *sal^E/Pv^>*). Wgn accumulated in the cells surrounding the apoptotic zone (Fig. 5A). Contrastingly, apical Grnd localization in the tissue surrounding the apoptotic zone did not vary (Fig. 5B). Wgn and Grnd were also detected in cellular debris in the *rpr-*apoptotic zone. Moreover, we also reduced ROS production in the *rpr*-ablated region (*sal^E/Pv^>rpr*, *Sod1*:*Cat*) and found a decrease in Wgn, but not Grnd, levels in the cells surrounding the ablated zone (Fig. 5C, 5D).

**Figure 5.**
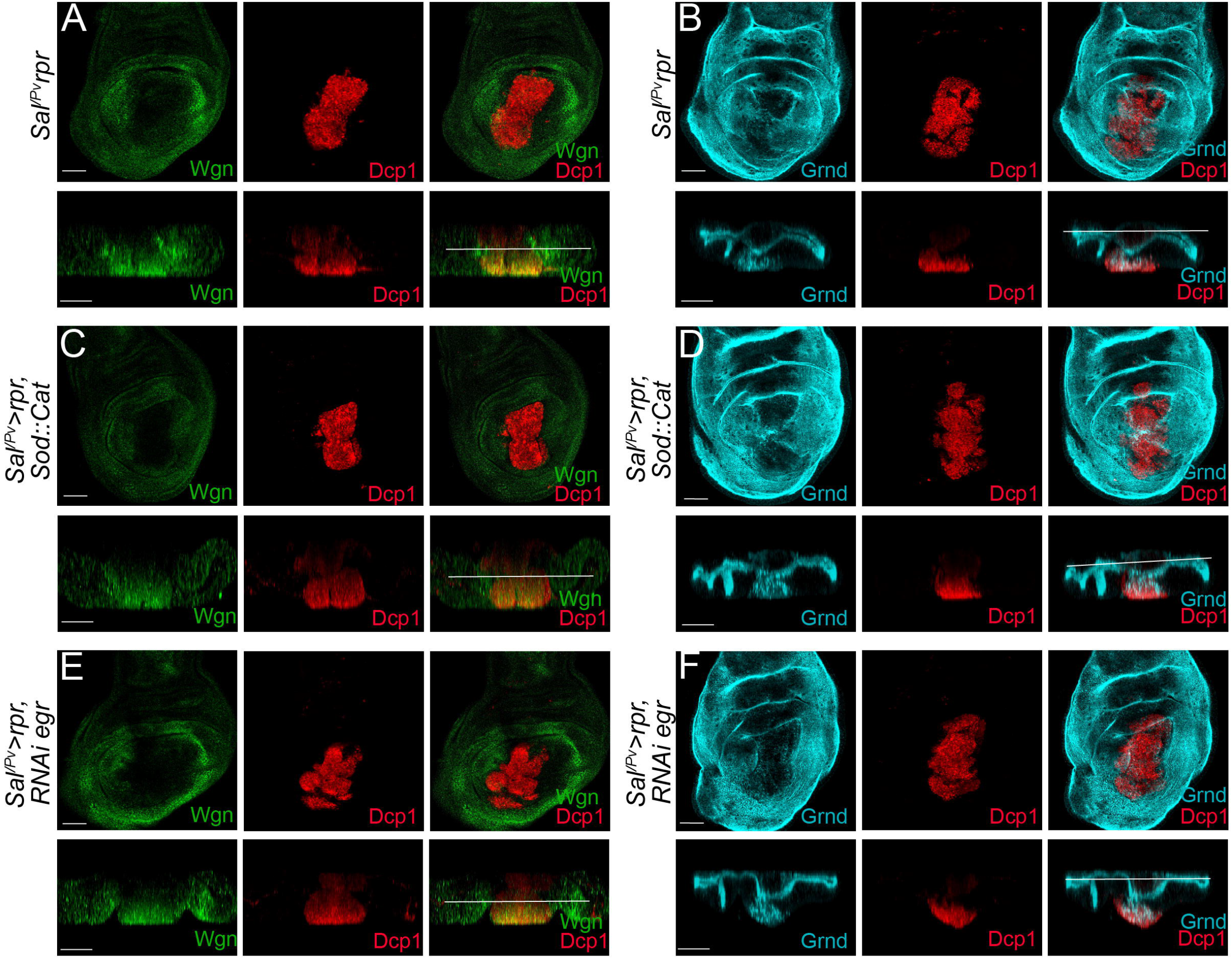
Wgn accumulation after ectopic activation of the pro-apoptotic gene *rpr* is abrogated by enzymatic depletion of ROS, independently of Egr/TNFα. The wing pouch of each genotype (above) is complemented with an orthogonal apico-basal section (below); white lines indicate the level of the z-axis of the above images. (A) Anti-Wgn and Dcp1 of disc with genetically induced apoptosis using s*al^E/Pv^>rpr* (n=12). (B) Anti-Grnd and Dcp1 of disc with genetically induced apoptosis using s*al^E/Pv^>rpr* (n=10). (C) Anti-Wgn and Dcp1 of s*al^E/Pv^>rpr, Sod1*:*Cat* (n=16). (D) Anti-Grnd and Dcp1 of s*al^E/Pv^>rpr, Sod1*:*Cat* (n=8). (E) Anti-Wgn and Dcp1 of s*al^E/Pv^>rpr, RNAi egr* (n=19). (F) Anti-Grnd and Dcp1 of s*al^E/Pv^>rpr, RNAi egr* (n=10). Scale bars: 50µm.

In addition, we found that *egr/TNF*α is autonomously expressed after inducing apoptosis by *rpr* (Fig. S3A-B). To test if the effects on Wgn localization were due to ROS or to the expression of *egr*, we used RNAi to knock down *egr* in the apoptotic cells and found that reduced Egr/TNFα had no effect on Wgn localization (Fig. 5E, 5F).

Together, these observations suggest that Wgn responds to apoptotic production of ROS independently of Egr/TNFα.

### Wgn, but not Grnd, is necessary for p38-dependent regeneration

Regeneration in the gut and in imaginal discs depends on p38 in a ROS-dependent manner (Patel et al., 2019; Santabárbara-Ruiz et al., 2019). Therefore, we wondered whether Wgn is necessary for p38-dependent regeneration. We used a double transactivation system to simultaneously induce apoptosis in one domain of the wing disc to stimulate regeneration and to knock down either *wgn* or *grnd* in adjacent regenerating cells (Fig. 6A). We first confirmed that the mutants for *wgn* and *grnd* used in this work do not affect normal growth and patterns (Fig. S4). However, knocking down *wgn* after inducing apoptosis resulted in anomalous wings, a characteristic of incomplete regeneration (Fig. 6B). Most of these wings showed reduced size and a defective pattern of veins and interveins. In contrast, wings carrying a knock down of *grnd* did not show a reduced size or defects in wing patterning (Fig. 6B), suggesting that regeneration of the apoptotic zone was completed even in the absence of *grnd.* To confirm this observation, we induced cell death (s*al^E/Pv^>rpr*) in an independent *grnd* mutant, *grnd^minos^*, in homozygosis and heterozygosis and found that wings regenerated normally. This result demonstrates that *wgn*, but not *grnd*, is necessary for the regenerative response after apoptosis.

**Figure 6.**
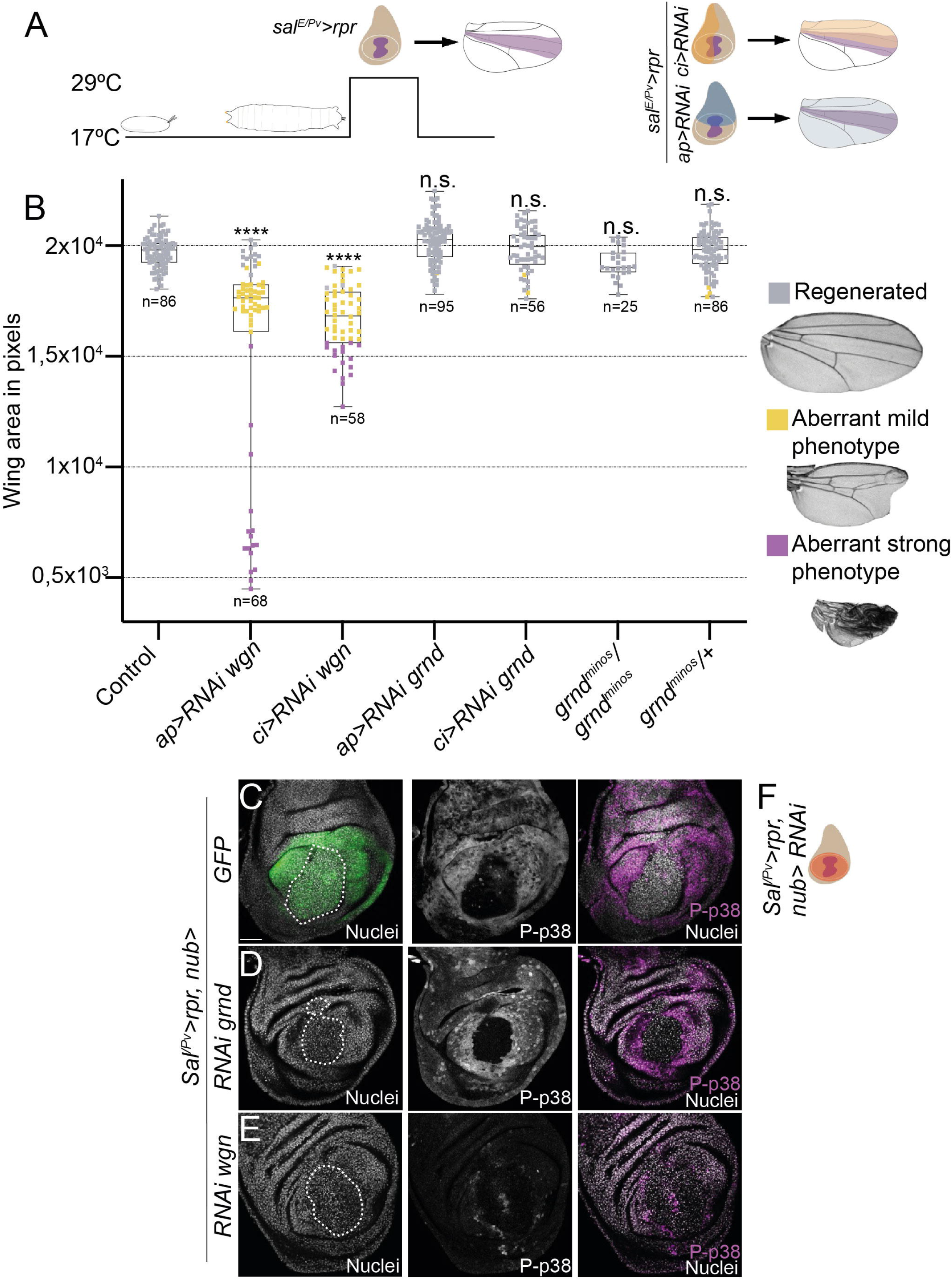
Knockdown of *wgn* but not *grnd* impairs p38-dependent regeneration. (A) Design for ectopic expression of transgenes and simultaneous apoptosis induction. Purple area in disc: apoptosis induced by s*al^E/Pv-^LHG,LexO-rpr*. Light purple in adult wing: regenerated tissue. The *Gal4/UAS* activates *RNAi* in *ci*/anterior compartment (yellow), *ap*/dorsal compartment (blue). Adult wings were scored for complete regeneration of the missing zone. s*al^E/Pv-^LHG* and *Gal4* are under the control of *tub-Gal80^TS^*. Apoptosis was induced by shifting the temperature to 29 °C for 11 hours. The larvae were transferred to 17 °C where they regenerated and emerged into adults, in which wings were scored. (B) Regeneration assay. Box plot: Y-axis shows the average area in pixels of adult wings obtained after apoptosis in the s*al^E/Pv^* region and the *RNAi* or mutant (genotype, X-axis). Each dot represents one wing; wild-type pattern/regenerated (gray), mild aberrant phenotype (yellow) and strong aberrant phenotype (purple), n.s.= not significant, \*\*\*\**p*<0.0001. (C-E) Activated p38 in regeneration assay of wing discs. Phosphorylated p38 (P-p38) of discs after s*al^E/Pv-^LHG,LexO-rpr* apoptosis and *nub-Gal* activation of *wgn* or *grnd UAS-RNAi.* (C) GFP (n=8), (D) *RNAi grnd* (n=18), and (E) *RNAi wgn* (n=12). White lines in the confocal images outline the pyknotic nuclei of apoptotic cells. (F) Design for ectopic expression of transgenes and simultaneous apoptosis induction when moved to 29 °C. Purple area: s*al^E/Pv-^ LHG, LexO-rpr* and light orange area corresponds to *nub>RNAi.* TP3 used to stain the nuclei. Scale bar: 50µm.

Next, to test if *wgn* is required for p38-driven regeneration, we analyzed if phosphorylated p38 is affected by *wgn* downregulation after inducing apoptosis. We used the double transactivation system to induce apoptosis in the *Sal*-central zone of the wing pouch and to knock down *wgn* or *grnd* in the wing pouch (henceforth *nub>*) (Fig. 6F). In control apoptotic discs, we detected activation of p38 in the wing pouch surrounding the apoptotic cells (Fig. 6C), as previously described (Esteban-Collado et al., 2021; Santabárbara-Ruiz et al., 2015, 2019). Likewise, an activation of p38 in cells near the apoptotic zone was found after knocking down *grnd* (Fig. 6D). However, after knocking down *wgn*, phosphorylated p38 in the wing pouch surrounding the apoptotic cells was abolished (Fig. 6E). These results demonstrate that *wgn*, but not *grnd,* is key for the activation of p38 signaling during the regenerative response to apoptosis.

## Discussion

In this work we have demonstrated that the conserved TNFR Wgn is activated by oxidative stress, accumulates near the source of ROS, and confers survival to cells in the absence of Egr/TNFα. This function is primarily focused on exacerbating p38 activity, likely via dTRAF1/Ask1, and is tightly linked to the oxidative stress released after cell damage.

Mammalian TNFR pathways have pleiotropic functions as a result of complex regulatory mechanisms (Gough & Myles, 2020). Upon binding to TNFα, a core signaling complex is constructed on the cytoplasmic tail of the TNFRs. This signaling complex includes the TRAF adaptor proteins as major signal transducers for the TNFRs. In mammals and *Drosophila,* a range of biological functions, such as adaptive and innate immunity, embryonic development, and stress response, are mediated by TNFRs/TRAFs through the induction of cell survival, proliferation, differentiation or cell death. Thus, TRAFs add complexity to the upstream TNFR and consequently to the signal transduction (Chung et al., 2002). In addition to TNFR, TRAFs associate to MAP3 kinases such as Ask1 or Tak1, which will trigger the p38 or JNK MAP kinase pathways (Hoeflich et al., 1999; Nishitoh et al., 1998). In mammals, once TNFR1 is activated and binding has occurred with the core complex, typically comprised of TRAF2, TRAF5 and the receptor-interacting kinase RIPK1, post-translational modifications will determine the ability to activate p38 or JNK for survival and inflammation (Dostert et al., 2019). If these protein modifications are disrupted, the signaling complex triggers apoptosis (Dostert et al., 2019; Vince et al., 2009). The *Drosophila* Wgn signaling is also pleiotropic and its function in the cell will also depend on upstream and downstream signals. Wgn was first described as pro-apoptotic after binding to Egr/TNFα (Kanda et al., 2002), with dTRAF2 or dTRAF1 interacting with the TNFR (Geuking et al., 2005; Kauppila et al., 2003). However, the lack of the death domain, distinctive of many TNFRs, and the inability of *wgn* mutants to rescue the apoptotic phenotype generated by Egr/TNFα, suggests that apoptosis is not the main function of *wgn* (Andersen et al., 2015; Kanda et al., 2002). Indeed, Wgn is emerging as a pro-survival TNFR and, as in mammals, the combination of TRAFs or other adapter proteins in the C-terminal core complex could divert the pathway towards functions other than apoptosis. Indeed, our regeneration assay, where *Drosophila* Wgn-dTRAF1 has a survival function, as well as where Wgn-dTRAF3 suppresses lipolysis in the gut, are examples of a *Drosophila* TNFR acting independently of Egr/TNFα (Loudhaief et al., 2023). In contrast, Grnd/dTRAF2 ensures the pathway towards JNK-driven apoptosis in a ligand-dependent manner, likely through the Tak1 MAP3 kinase (Andersen et al., 2015; Palmerini et al., 2021). Furthermore, Grnd displays nanomolar binding affinity for Egr that is three orders of magnitude higher than it is for Wgn, suggesting that canonical Egr signaling in *Drosophila* occurs predominantly through Grnd activation rather than through Wgn (Palmerini et al., 2021).

The TNFR superfamily has been described to contain a CRD in their N-terminal end. Cysteine residues have active thiols that can be efficiently oxidized by ROS, a physiological mechanism to transmit external signals to the intracellular system (Gotoh & Cooper, 1998; Kamata et al., 2005; Zhang et al., 2001) and subsequently alter protein structure, interaction with partners, and subcellular localization (Sies et al., 2017). Moreover, human TNFR1/2 can suffer oxidative stress-induced self-association due to modifications of the CRD resulting in ligand-independent signaling (Ozsoy et al., 2008). Therefore, we speculate that as in mammals, the oxidation of Wgn could promote its self-association and influence the partner preferences in the core signaling complex, i.e., Wgn-dTRAF1, in a ligand-independent manner. In addition, dTRAF1 has been described to be a positive interaction partner of Ask1 (Kuranaga et al., 2002). Hence, we propose that Wgn-dTRAF1-Ask1 is a signaling module activated upon oxidative stress to ensure survival in cells that will be involved in the regenerative response. Moreover, we recently demonstrated that Ask1 requires the activity of the nutrient-dependent Pi3K/Akt signal to divert Ask1 function to p38 phosphorylation in cell survival and regeneration (Esteban-Collado et al., 2021). Thus, we conclude that it is not only Pi3K/Akt, but also Wgn, that will be necessary for leading Ask1 to induce a p38 response to apoptosis.

Here, we propose a model for the targets that respond to ROS upon cell damage. Cells that have been damaged, either by injuries or apoptosis, produce ROS, normally of mitochondrial origin (Murphy, 2009). ROS spreads from damaged or dying cells to the nearby healthy cells, acting as early signals for tissue recovery in different organisms such as flatworms and mammals (Gauron et al., 2013; Rampon et al., 2018). In our model, the ROS-dependent post-translational modifications will primarily target three branches that converge with Ask1 (Fig. 7): (a) Wgn, which will recruit dTRAF1 to the Wgn core signaling complex, (b) the thioredoxin bound to the inactive Ask1 signaling complex, which upon oxidation will be dissociated and allow the active Ask1 to interact with dTRAF1 (Sakauchi et al., 2017), and (c) the insulin/Pi3K/Akt signaling pathway that will phosphorylate Ask1 for survival and phosphorylation of p38 (Esteban-Collado et al., 2021; Santabárbara-Ruiz et al., 2019). Thus, the divergent role of Grnd and Wgn is driven not only by the different affinity for the ligand Egr/TNFα, but also by the ROS-dependent activation of Wgn. We have shown here that Wgn accumulation near the damaged zone can be reverted after ROS depletion, indicating that ROS produced by dying cells is involved in the Wgn response after damage. Lineage experiments showed that these cells near the damaged zone are responsible for most of the regenerated epithelium (Bosch et al., 2008; Repiso et al., 2013). However, we cannot rule out that the accumulation of Wgn near the wounded area responds not only to a reorganization of vesicles, but also to a transcriptional response. Indeed, RNAseq of regenerating imaginal discs has shown that *wgn* is transcriptionally upregulated in the earliest phase of regeneration (Vizcaya-Molina et al., 2018).

**Figure 7.**
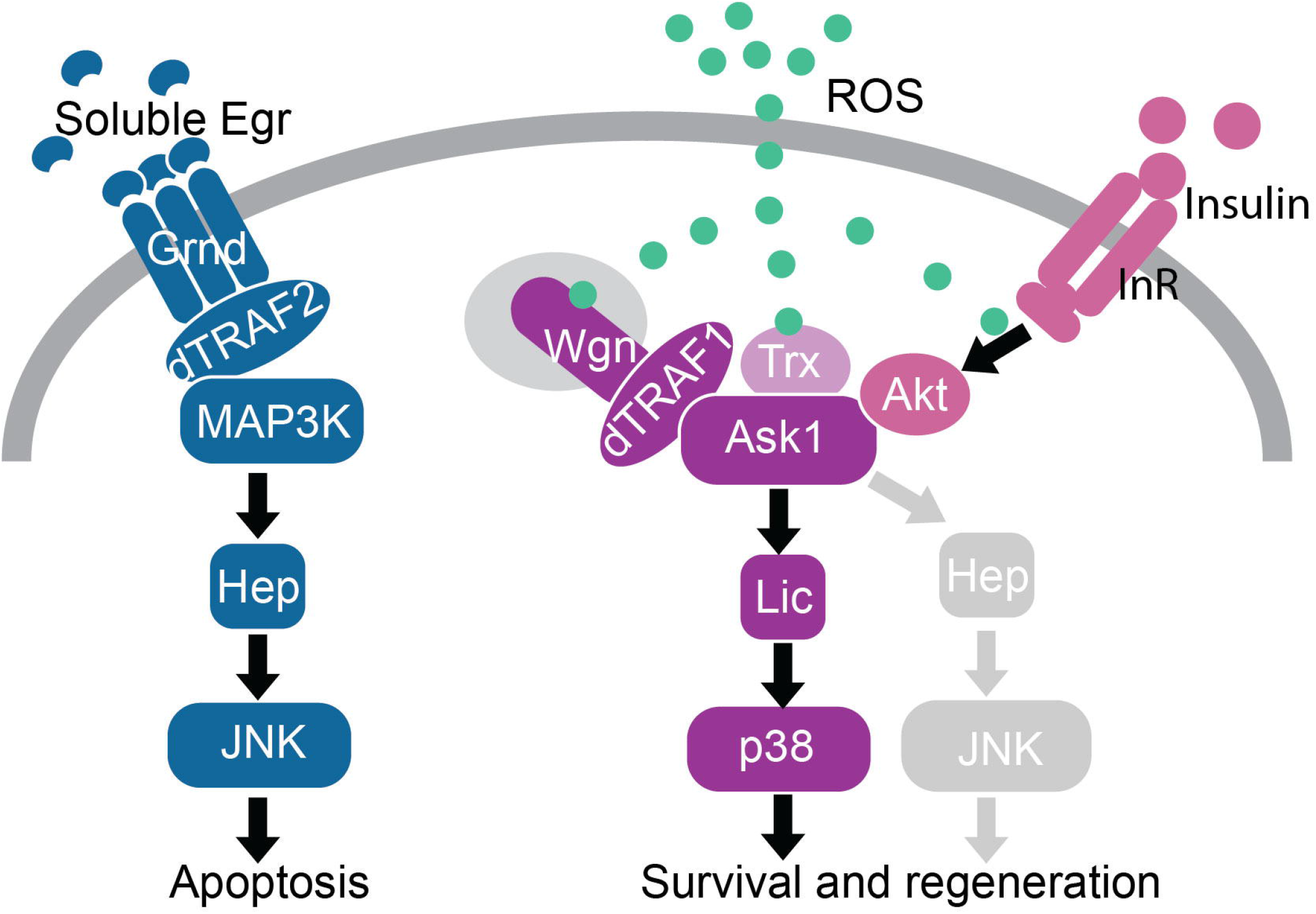
Model for ROS-dependent Wengen recruitment for survival and regeneration. ROS are required for the oxidation of thioredoxin (Trx) to dissociate from Ask1. ROS are required for the phosphorylation of Akt, which in turn phosphorylates Ask1 to divert its function towards survival and regeneration. ROS are required for Wengen recruitment in the Ask1/p38 axis. Phosphorylation of Ask1 by Akt results in tolerable levels of p38 and JNK (perhaps very low, gray) activity and the avoidance of apoptosis.

We observed a very high degree of divergence between Grnd and Wgn, showing that both gene families are of extremely ancient origin and are found across pancrustacean and arthropod lineages, respectively. This taxonomic distribution suggests that Grnd originated through a duplication and fast divergence at the base of pancrustaceans. These results match the subfuncionalization observed of both genes: Grnd having a pro-apoptotic function and Wgn promoting cell survival. The amplification and subfunctionalization of TNFRs has also been observed in lineages for adaptation to biotic or abiotic stress (Quistad & Traylor-Knowles, 2016).

Remarkably, we observed that there is also a very high sequence divergence between the CRDs of Grnd and those of all the other TNFR families we studied. By contrast, Wgn CRDs were much more canonical, and we found that they clustered together with the CRDs from deuterostome TNFRs, including those from humans.

We speculate that the subcellular location of Wgn and Grnd may contribute to the different functions of both receptors. Grnd is more exposed at the apical side of the plasma membrane, which makes this receptor more accessible for ligand interactions (Agrawal et al., 2016). Wgn, embedded in cytoplasmic vesicles, is less accessible to the ligand and could be more restricted to being activated by local sources of signaling molecules, such as ROS.

## Acknowledgements

The authors would like to thank Dr. Manel Bosch from the Optical Microscopy Unit of the CCiT for assistance. This research was funded by the Spanish Ministerio de Ciencia, Innovación y Universidades (PID2021-123300NB-100) and by the Agència de Gestió d’Ajuts Universitaris i de Recerca (2021SGR00293).

## Materials and methods

### Drosophila strains

Animals were reared on standard fly food. The *Sal^E/Pv^ -LHG* and *lexO-rpr Drosophila* strains have been previously described (Santabárbara-Ruiz et al., 2015). Other strains were: *UAS-egr^weak^* (Moreno et al., 2002), *UAS-lic^wt1.1^*(Terriente-Félix et al., 2017), *UAS-hep^wt^* (Uhlirova & Bohmann, 2006), *Sal^E/Pv^-Gal4* (Barrio & de Celis, 2004), *ptc-Gal4* (Hinz et al., 1994), *tub-Gal80^TS^* (McGuire et al., 2003), c*i-Gal4* (Martin & Morata, 2006). The *eiger-lacZ* was provided by K. Basler. From the Bloomington Drosophila Stock Center (BDSC): *UAS-rpr* (BL92781), *UAS-Sod1.a* (BL24754), *UAS-Cat.a* (BL24621), *UAS-p35* (BL5072), *UAS-RNAi wgn* (BL55275), *UAS-RNAi Ask1* (BL35331), *UAS-GFP* (BL4776), *tub-Gal80^ts^* (BL7017) *UAS-RNAi Tak1* (BL53377). From the Vienna *Drosophila* Resource Center (VDRC): *Egr-GFP* (Vienna 318615), *UAS-RNAi dTRAF1* (KK110766), *UAS-RNAi grnd* (KK104538), *UAS-RNAi wgn* (KK108814), *UAS-RNAi dTRAF2* (KK110266). *UAS-RNAi eiger* (Kuranaga et al., 2002). The *w^118^;+;+* strain was used as a control.

### Immunolocalization

The primary antibodies used in this work were against phospho-p38 (rabbit 1:50, Cell Signaling Technology, 9211S), cleaved *Drosophila* caspase 1 (Dcp1) (rabbit 1:200, Cell signaling, 9578S), ß-galactosidase (mouse 1:1000, Z3783), Wengen (mouse 1:100, gift from K. Basler), Grindelwald (guinea pig 1:500, gift from Pierre Leopold). The fluorescently labeled secondary antibodies were from Life Technologies, all used 1:200 in 0.3% PBT: goat anti-mouse Alexa Fluor 488, donkey anti-rabbit Alexa Fluor 532, and goat anti-guinea pig Alexa Fluor 555.

Wing imaginal discs dissected from late third instar larvae in 1×PBS (pH 7.4) were fixed in 4% paraformaldehyde (Electron Microscopy Sciences) in PBS for 40[minutes at room temperature, washed in PBS 0.3% Triton X-100 (PBT), blocked for 2 hours in PBT containing 2% BSA, and incubated overnight with primary antibodies at 4 °C. The next day, the discs were washed, and incubated with secondary antibodies for 2 hours at room temperature. After washing, nucleic acid staining was performed by incubating the discs for 10-15 minutes with the nuclear marker TO-PRO-3, 1mM (TP-3, Life Technologies) or 10 mM of DAPI (Life Technologies). The discs were mounted in SlowFade Diamond Antifade Mountant (Life Technologies).

### Image acquisition

For the confocal images, a Zeiss LSM880 and a Leica SPE confocal laser scanning microscopes were used. Images were analyzed and processed using FIJI. A Leica DMBL microscope was used for taking pictures of the adult wings.

### Statistical analysis and cell death area ratio calculation

To calculate the cell death area ratio, we used FIJI. First, we generated a Z-projection of the whole stack for the anti-Dcp1 channel using the Max intensity projection. Then, we applied the remove outliers tool (radius 2, threshold 10). Subsequently we thresholded the image with the MaxEntropy Threshold to create a binary image and automatically determine the area in pixels of the dead zone with the analyze particles tool. The data was normalized dividing the area of Dcp1-positive cells by the total area of the imaginal disc. As a result, the cell death area ratio was obtained.

In Figures 1 and 3, the data are in mean ± SD. To make statistical comparisons, we used a one-way analysis of variance (ANOVA) followed by Tukey’s post-hoc test to make pair comparisons between each group using GraphPad. Significance is indicated in the figures only when *p<*0.05, as follows: *\*p<*0.05, *\*\*p<*0.01, *\*\*\*p<*0.001, and *\*\*\*\*p<*0.0001.

### Gal4/UAS/Gal80ts for *eiger^weak^* activation in the wing imaginal disc

The Gal4/UAS transactivation system was temporarily controlled by the *tub-Gal80^TS^* thermo-sensitive *Gal4-*repressor. *Egr/TNF*α was ectopically expressed using the *UAS-eiger^weak^* construct (Moreno, Yan, et al., 2002), which results in reduced activity of *egr* and therefore low levels of apoptosis. The expression of the transgene was controlled by the thermo-sensitive Gal4 repressor *tub-Gal80^TS^*.

*Drosophila* of the desired genotype were cultured to lay eggs for 24 hours at 17 °C. Conditions for experiments in Figures 1 and 3: Embryos were kept at 17 °C until the 8th day (192 hours) after egg laying to prevent *UAS-eiger^weak^* expression. The larvae were subsequently transferred to 29 °C for 16 hours and then the imaginal discs from wandering larvae were dissected and processed for staining and immunofluorescence studies.

Conditions for experiments in Figure 4: to enhance the production of *egr*-dependent apoptotic cells, embryos were kept at 17 °C until the 7th day (168 hours) after egg laying to prevent *UAS-eiger^weak^*expression. The larvae were subsequently transferred to 29 °C for 24 hours and then the imaginal discs from wandering larvae were dissected and processed for staining and immunofluorescence studies.

### ROS detection *ex vivo*

All experiments for ROS detection were done in living conditions. To detect the presence of ROS, we used the MitoSOX reagent (Life Technologies), which is an indicator of oxidative stress of mitochondrial origin in living cells. Third instar discs were dissected in Schneider’s medium immediately after cell death or injury and incubated in agitation for 15 minutes in medium containing 5 μM MitoSOX reagent, followed by three washes in Schneider’s culture medium (Sigma-Aldrich S0146). The samples were protected from light throughout the experiment. They were then mounted using culture medium supplemented with 1 μM TO-PRO-3 (Life Technologies) for nucleic acid staining. As *in vivo* TO-PRO-3 only enters dying and dead cells, so we used it to distinguish dead cells from living cells.

### Genetic ablation and the dual Gal4/LexA transactivation system

For adult wing regeneration analysis, we used a dual Gal4 and lexA transactivation system, as described previously (Esteban-Collado et al., 2021; Santabárbara-Ruiz et al., 2019). The *UAS-RNAi wgn* and *UAS-RNAi grnd* transgenes were activated in *ci/ap* domains of the wing disc using the Gal4/UAS system. Apoptosis was induced during larval stages in the wing-specific *sal^E/Pv^* domain using the *Gal80-*repressible transactivator system LHG (LexA-Hinge-Gal4 activation domain), a modified form of the *lexA/lexO* system (*-LHG>lexO-rpr*) (Yagi et al., 2010) (Fig. 6A). After the adults emerged, the wings were dissected and regeneration was analyzed.

For testing the capacity to regenerate, we used adult wings from *sal^E/Pv^>rpr* individuals, in which patterning defects can be easily scored. Flies were fixed in glycerol:ethanol (1:2) for 24 hours. Wings were dissected in water and then washed with ethanol. They were then mounted in 6:5 lactic acid:ethanol and analyzed and imaged under a Leica microscope.

The areas of the mounted wings were outlined and scored using FIJI. In addition, we divided wings in three categories depending on the strength of defects in terms of pattern and number of absent veins and interveins: regenerated (normal pattern), mild aberrant phenotype (1-2 veins missing) and strong aberrant phenotype (>3 veins missing or more aberrant phenotype).

The use of the knock down RNAi transgenes alone did not affect vein pattern or wing size (Fig. S4). Also, flies carrying the constructs shown in Figure 6 but grown constantly at 17 °C to maintain *tub-Gal80^TS^* activity and block transgene (*UAS or lexO*) expression, did not show defects in the wing size or pattern (Fig. S4)

### Characterization of Grnd and Wgn gene family evolution

Protein sequences of TNFRs were researched using the software blastp and tblastn from the online Blast server (https://blast.ncbi.nlm.nih.gov/Blast.cgi), restricting by taxonomy to specifically search in the genomes and proteomes of those animal lineages of particular relevance for Grnd and Wgn evolution. *Drosophila melanogaster* Grnd and Wgn were used as initial queries. After that initial search, a second one was performed using the retrieved protein hits as new queries. The same procedure was used in two slow-evolving deuterostome lineages (a cephalochordate, the European amphioxus *Branchiostoma lanceolatum*, and a hemichordate, the acorn worm *Saccoglossus kovalevskii*), using *Homo sapiens* TNFRs as initial queries.

For the phylogenetic analyses of TNFRs, protein sequences were aligned with the orMAFFT software (Katoh et al., 2019) and the resulting alignments were trimmed using Aliview to discard spuriously aligned regions (Larsson, 2014). Phylogenetic trees were built using IQ-Tree (Nguyen et al., 2015), testing the tree with UFBoot (bootstrap = 103) (Hoang et al., 2018) and an approximate Bayes test for single branch testing. The model used was selected using ModelFinder with BIC as the criteria (Kalyaanamoorthy et al., 2017). The trees were visualized using ITOL (Letunic & Bork, 2021).

Two phylogenetic trees were built, one using only the CRDs from all the arthropod and deuterostome TNFRs retrieved in our searches and a second one using the full protein sequences of Wgn and Grnd from different arthropods, except a few highly divergent sequences from Ostracoda and Thecostraca.

For the CRD tree, the corresponding protein domains were extracted using Hmmer 3.4 (http://hmmer.org/). LIM domains, which are also cysteine-rich domains unrelated to those of TNFRs, from the human proteins ISL1, LIMCH1, and FHL1 were used as an outgroup. The model selected for this tree was PMB+G4.

The Grnd and Wgn tree was built with the model VT+I+G4+F. Three TNFRs from deuterostomes were used as outgroups, TNFR6L from *B. lanceolatum* (CAH1240663.1), TNFR11L from *S. kowalevskii* (XP_006819214.1), and TNFR1B from *H. sapiens*.

A PCA-based alignment of all the CRDs used in the previous phylogenetic analyses was built using the Jalview software (Waterhouse et al., 2009) and represented with R. Also, an all versus all comparison was performed using blastp. The percentage of identity was normalized with the total length of the query. The results were represented by R (github.com/rlbarter/superheat).

---

**Supplementary figure 1.** Fig. S1. (A-B) Dcp1-positive cells after *egr^weak^* overexpression in (A) *RNAi wgn* KK strain and (B) *RNAi wgn* TRIP strain (see materials and methods section). TP3 used to stain nuclei. Scale bar: 50µm.

**Supplementary figure 2.** Fig. S2. (A) Full version, including all the species names and sequence accession numbers of the ML phylogenetic tree of the CRDs of Wgn, Grnd, and deuterostome TNFRs featured in Fig. 2A. *B. lanceolatum*, *S. kowalevskii,* and *H. sapiens* are abbreviated as Bla, Sko and Hsa, respectively. (B) ML phylogenetic tree of all identified arthropod proteins, using full-length protein sequences and three deuterostome TNFRs as outgroups, *B. lanceolatum* TNFR6L (CAH1240663.1), *S. kowalevskii* TNFR11L (XP_006819214.1), and *H. sapiens* TNFR1B. (C) Pairwise comparison of Wgn, Grnd, and deuterostome CRDs. (D) CRD alignments of some representative Wgn and Grnd proteins.

**Supplementary figure 3.** Fig. S3. (A-B) Apoptosis genetically induced in *ptc>rpr* in two different Egr reporter backgrounds; (A) *Egr2xGFP* reporter and (B) *EgrLacZ* detected by anti-ß-Gal antibody. The yellow zone in the merged image shows colocalization of ß-Gal-positive cells and Dcp1-positive cells (dead cells). The dotted lines outline pyknotic nuclei of apoptotic cells. TP3 used to stain the nuclei. Scale bars: 50µm.

**Supplementary figure 4.** Fig. S4. Control for regeneration assay. (A) Box plot: Y-axis shows the average area in pixels of adult wings obtained from controls kept at 17 °C, with no cell death induction (s*al^E/Pv-^LHG,LexO-rpr* OFF) and no expression of the transgenes. It also shows the average area in pixels from adult wings after the sole expression of the *RNAi* or mutant background (genotypes indicated in the X-axis). Each dot represents one wing: wild-type pattern (gray).

**Supplementary table 1.** Wgn and Grnd gene complements in arthropods.

