## Supplementary material for "ROS-mediated TNFR Wengen activation in response to apoptosis": Suppl Figures and Table

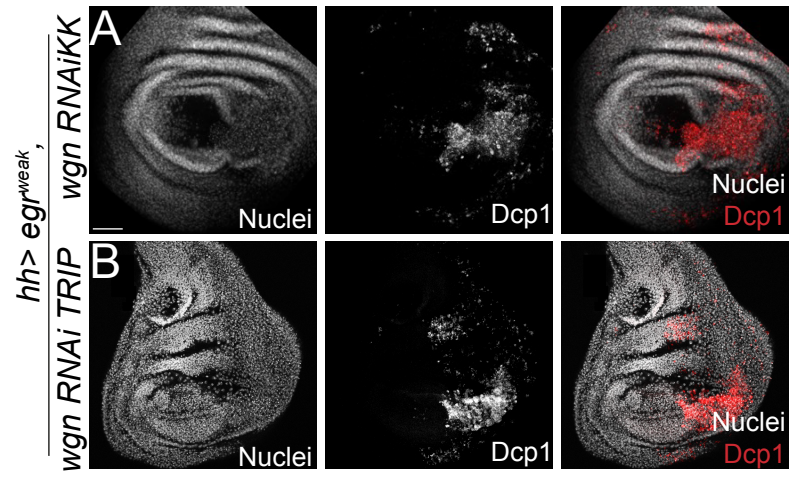

Figure supplementary 1

A

Branch support

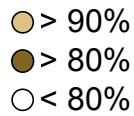

B

Branch support

● >90%
● >70%

D

LOC129908656-Wgn-[Episyrphus balteatus]/56-101
Wgn-[Drosophila melanogaster]/87-132
XP\_037968220.1-Wgn-[Plutella xylostella]/28-76
LOC664119-Wgn-[Tribolium castaneum]/30-75
LOC100651623-Wgn-[Bombus terrestris]/33-78
LOC110832081-Wgn-[Zootermopsis nevadensis]/28-75
LOC106669879-Wgn-[Cimex lectularius]/20-65
CAB3373027.1-Wgn-[Cloeon dipterum]/21-67
LOC124198684-Wgn-[Daphnia pulex]/53-99
LOC111711857-Wgn-[Eurytemora carolleeae]/50-95
LOC121871879-Wgn-[Homarus americanus]/26-72
LR903237-Wgn-[Darwinula stevensoni]/52-96
LOC129972442-Wgn-[Argiope bruennichi]/18-63

LOC129916031-Gmd-[Episyrphus balteatus]/14-62
Gmd-[Drosophila melanogaster]/35-83
CAG9138140.1-Gmd-[Plutella xylostella]/25-73
LOC660354-Gmd-[Tribolium castaneum]/31-79
LOC100644341-Gmd-[Bombus terrestris]/28-76
LOC110837103-Gmd-[Zootermopsis nevadensis]/10-58
LOC106667425-Gmd-[Cimex lectularius]/25-71
CAB3361506.1-Gmd-[Cloeon dipterum]/28-76
LJU01000149-Gmd-[Orchesella cincta]/21-68
LOC110852048-Gmd-[Folsomia candida]/27-74
LOC111707847-Gmd-[Eurytemora carolleeae]/10-55
FJT64021454-Gmd-[Amphibalanus amphitrite]/28-77
LOC121876844-Gmd-[Homarus americanus]/9-51

C

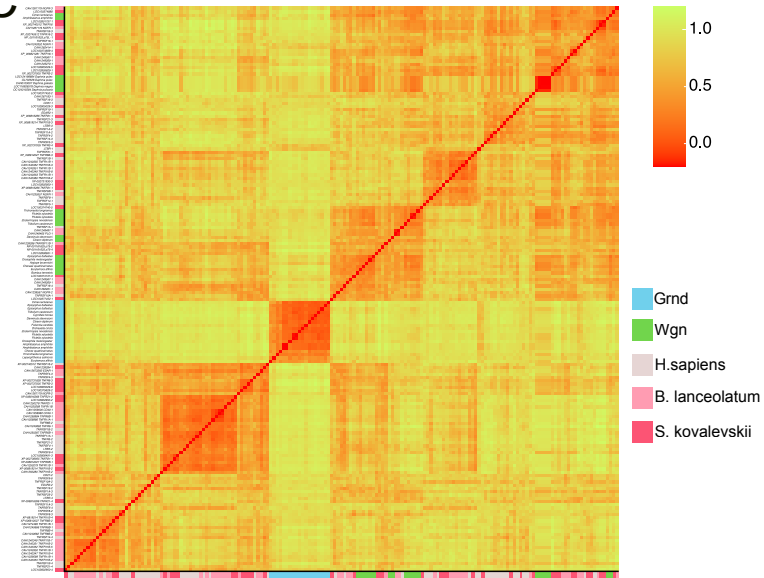

10 20 30 40 50
SPQ - Q SFQWWD P FQDK - - CIPCTEC - SGQKI - VLRPCQLHRTDI CGSIYDL
SPQ - APQHWWD SQDR - - CTPTRC - QGEM - PLRPCQLHTDI CGSIYDL
AVC ERGRSWDRRA - - CAPCSLCPARRLAV LRPCELHRTDVCALQHI
SIC - VSRQY FNSRLAS - - VVNOTEC - VEGDI - VVRPCQGHKDTVCGPMSNL
PVCKPGFEFWSEHAT - - CWPCTRC - APEF - T LSPCAVYKDAICGPLSAL
ALORPGHGFWD ETSER - - CVNCSKCPVAGHV - VLRPCQVHQDTQCGPLSAL
RLQ - NLNEYWSNEEI - - CLPOTKC - PPHQL - TMRPCQGHKDTVCGPMSNL
PPQPKWGHGFWDNRQE - - CAQCKVC - SHNEV - VLMPCQPYKDTL CGPMSAL
DFCQGVWAFKDPOTGR - - CLPQSPC - PEDHL - TVAKCFEDRTLCRPLTDL
- - CRRVFY FWD TGK - - CLPOTRCQTRGLV - TLLPCQYFRDSVCVKSSDW
LSCQQGQEFYNS EQGR - - CVPCTRC - HGSLV - VVVPCYVYRDAV CAPAAQF
HACI EGTYSQD - - TSEILDCFLSGC - GDGLF - ELHACNPFRTDICDSC - -
ELQKQKEWFND - MTGK - - CQPCITVC - AGKSW - EMAPCLDFSDTLVNLI EL

10 20 30 40 50
CNN - AKVCSL - EEFYCTSFSECEKEEIDTISHNFHESDCKNYGPAYGND
CHG - TICHVP - NEFCYVATERCHP - IEV - CNNQTHNYDAFLCAKECSAYKTF
KCG - QLKCAA - DDYCSPTDRCAP - CSDVCTKGHNFDGLVKECQGYIHN
SCG - QKKCKR - DEYCTSYKACEP - CSKICDKSTHNFDETCEVKKCQDYIHD
DCG - EQRCS - VEYCSYDKRCKP - CSSI - CDATGRNYQQNECICQCEYLHD
VCG - QKTKCT - KEYCSNDHMMCR - PDEVCETSNNYQKQTCEDQCVYIYD
PCG - EKSCSR - YECSNDLY - - CONCEPCTVGERNFDSVCEKECEYELHD
LCG - VHTCSD - NOYNSFGNICSD - CSNICDLKSHNYEHQECFDKQGYIHD
- CG - EIVCKA - HYCARIDDS - CRQETCDRALSNYDELDCEKDCQDYLHD
- CG - DIICQP - HNYCSQIDHTCRFQVAC - DPTLGNYDKGVCEDQDWLHD
TCG - SDLPCTDLYS - CDTFLSRCP - CSQLC - - - KDNFSE - - CHQKCEGFLQS
RCG - TDHTCSL - HOYCSPATGT - CEAYQDICHPGTKNYGRDCEKDCQDYLHD
TCG - ALPTDA - NGSCSMIN - - - - - QQKCRVSSPFGPQRCD - DDCCTPISD

Figure supplementary 2

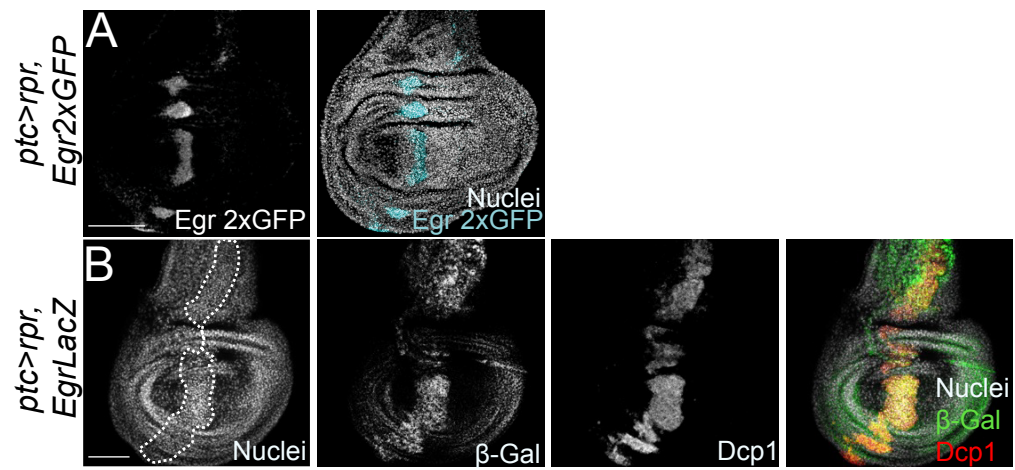

Figure supplementary 3

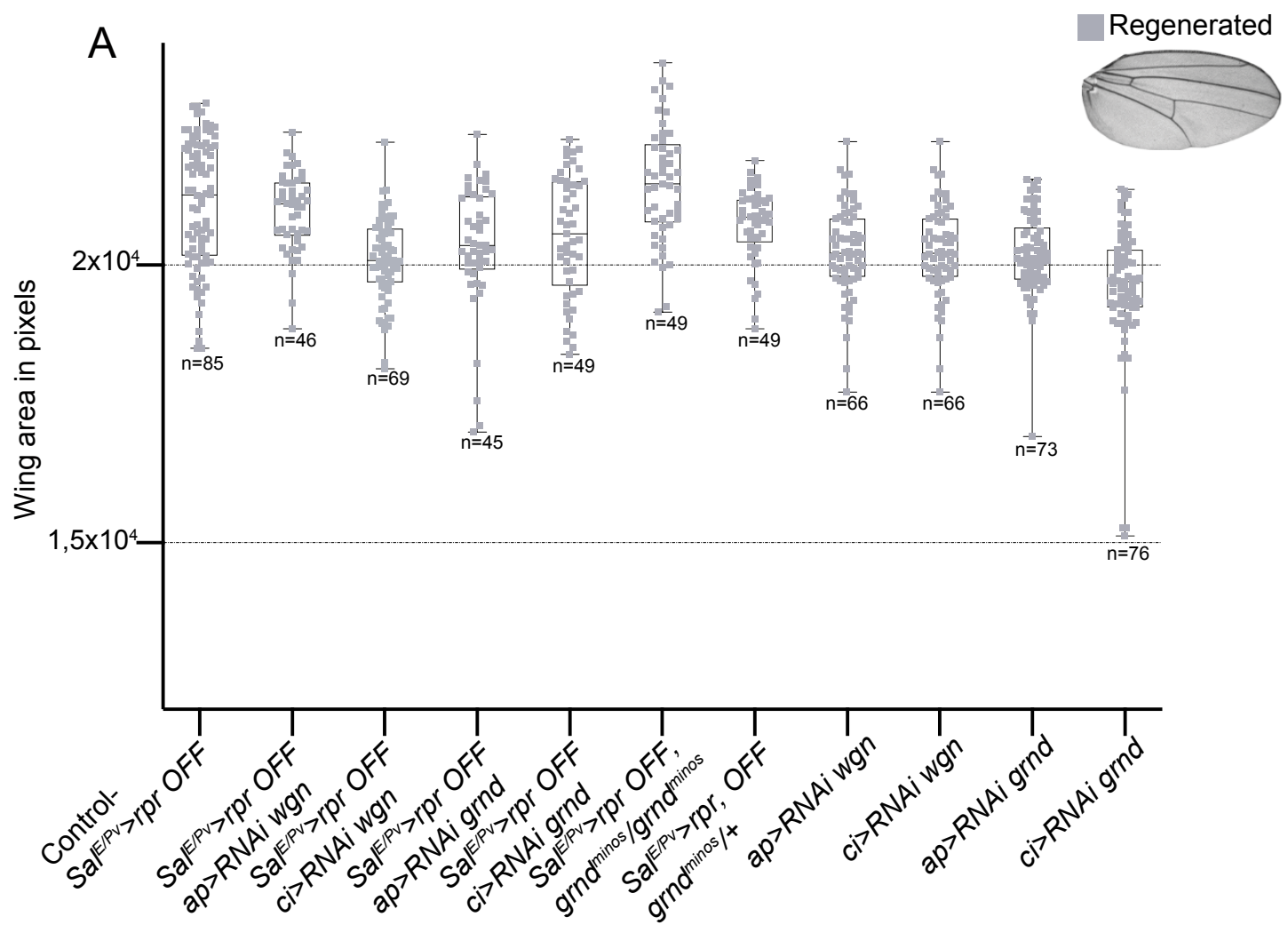

Figure supplementary 4

| Subphylum | Class/Superorder | Order | Sp | Grnd | Wgn |
| --- | --- | --- | --- | --- | --- |
| Chelicerata | Arachnopolmonata | Araneae | *Argiope bruennichi* | - | LOC129972442 |
|  |  | Scorpiones | *Hottentotta judaicus* | - | HQ288166 |
| Myriapoda | Chilopoda | Scutigermorpha | *Scutigera coleoptrata* | - | GCAQ01008710.1 |
|  |  | Lithobiomorpha | *Lithobius forficatus* | - | GCAY01031869.1 |
|  |  |  | *Eupolybothrus tridentinus* | - | GERX01040621.1 |
| Hexapoda | Insecta | Diptera | *Episyrphus balteatus* | LOC129916031 | LOC12990865 |
|  |  | Diptera | *Drosophila melanogaster* | CG10176 | CG6531 |
|  |  | Lepidoptera | *Plutella xylostella* | KAG7313426.1;CAG9138140.1 | KAG7305500.1; XP 037968220.1 |
|  |  | Coleoptera | *Tribolium castaneum* | LOC660354 | LOC664119 |
|  |  | Hymenoptera | *Bombus terrestris* | LOC100644341 | LOC100651623 |
|  |  | Hemiptera | *Cimex lectularius* | LOC106667425 | LOC106669879 |
|  |  | Blatodea | *Zootermopsis nevadensis* | LOC110837103 | LOC110832081 |
|  |  | Ephemeroptera | *Cloeon dipterum* | CAB3361506.1 | CAB3373027.1 |
|  | Collembola | Entomobryomorpha | *Folsomia candida* | LOC110852048 | - |
|  |  |  | *Orchesella cincta* | ODN01477 | - |
| Crustacea | Copepoda | Calanoida | *Eurytemora carolleeae* | LOC111707847 | LOC111711857 |
|  | Branchipoda | Diplostraca | *Daphnia pulex* | - | LOC124198684 |
|  | Thecostraca | Balanomorpha | *Amphibalanus amphitrite* | LOC122375700; FJT64_021454; LOC122381716 |  |
|  | Malacostraca | Decapoda | *Homarus americanus* | LOC121876844 | LOC121871879 |
|  | Ostracoda | Podocopida | *Cyprideis torosa* | CAD7232789.1 | - |
|  |  |  | *Darwinula stevensoni* | - | LR903237 |

Supplementary Table 1
